# A genetic tool targeting brain GPR75

**DOI:** 10.64898/2026.05.29.728898

**Authors:** Emma C. Wheeler, Runzhou Yang, Stephen M. Farmer, Sheng Zhang, Ningyan Zhang, Zhiqiang An, Qingchun Tong

**Author notes:** Correspondence: Qingchun Tong.

## Abstract

G-protein-coupled receptor 75 (GPR75) has emerged as an important mediator in diet-induced obesity (DIO) and a promising therapeutic target for anti-obesity drugs. However, the anatomical location of GPR75 in the brain remains unclear, hindering the understanding of GPR75 biology in DIO. Here, we generated a new GPR75-GFP-Ires-Cre knockin mouse strain, in which the Cre expression is driven by the endogenous GPR75 promoter and the GFP is fused with the C-terminal of the GPR75 protein. Both Cre and GFP were confirmed to be colocalized with the endogenous GPR75 expression. In addition, the GPR75-GFP fusion protein remains functionally normal with unaltered susceptibility to DIO. Moreover, using this mouse strain, we found that GPR75 is broadly expressed throughout the brain and mainly localized to the cytoplasm of brain neurons. This new genetic tool can therefore be used to study the neural basis for GPR75 in mediating DIO.

## Introduction

The current level of obesity in the global population has reached an alarming level and has consequently caused a huge burden to the patients and society. Despite intense research efforts, effective treatment against obesity has been difficult until recently, with the discovery of glucagon-like peptide-1 receptor (GLP-1R) agonism, which has become a cornerstone in the success of effective treatments of obesity ^1,2^. However, GLP-1R agonism-based treatments are known to have a high cost with adverse side effects, including nausea, and are not equally effective for all obesity patients, in addition to rebound weight gain with treatment withdrawal ^3,4^. Thus, alternative obesity treatment options are highly desired that can be used independently of or in combination with GLP-1R agonism to achieve more effective weight reduction.

G-protein-coupled receptor 75 (GPR75) has been shown to play an important role in diet-induced obesity (DIO) ^[5]^. Mutations in the GPR75 gene have been associated with reduced body weight in humans, and mice with total body GPR75 deletions (GPR75-KO) exhibit dramatic resistance to DIO ^5-7^ , suggesting a conserved role for GPR75 in mediating DIO in both rodents and humans. Recent studies suggest that the expression of GPR75 in glutamatergic, but not GABAergic, neurons mediate its action on DIO ^8^. Although previous studies suggest that 20-Hydroxyeicosatetraenoic acid (20-HETE) and C-C motif chemokine ligand 5 (CCL5)-function as endogenous ligands for GPR75 ^9^, recent results dispute these results ^10^, suggesting that GPR75 remains an orphan receptor. Importantly, despite its well-established expression pattern in peripheral tissues, the expression pattern of GPR75 in the brain is limited to *in situ* hybridization and *in vitro* cell culture ^10,11^. Thus, a genetic tool that specifically targets GPR75-expressing neurons is needed to examine the underlying biology for GPR75 in mediating DIO.

Here we report the generation and validation of a new GPR75-GFP-IRES-Cre mouse line, in which the Cre expression is driven by the endogenous GPR75 promoter, and GFP is fused in frame to the C-terminus of GPR75. This new mouse strain can be used to target GPR75-expressing neurons and examine the biology of GPR75 function.

## Results

Given the known importance of GPR75 in the brain for mediating DIO, and the lack of tools to target GPR75-expressing neurons and visualize GPR75, we aimed to generate a new mouse strain that can be used to target GPR75-expressing neurons and to track GPR75 protein location. Toward this, we generated a line of mouse that expresses Cre driven by the endogenous GPR75 promoter with the endogenous GPR75 replaced with GPR75-fusion protein (Fig. 1A). The correct DNA targeting was confirmed with the PCR genotyping using specific PCR primers for the wild-type and knockin alleles (Fig. 1B). To confirm whether GPF and Cre expression patterns follow that of GPR75, we performed in situ hybridization for GPR75 (green), EGFP (red), and Cre (blue). While the control C57 mice showed only GPR75 (Fig. 1C), GPR75-GFP-Ires-Cre mice exhibited signals for all 3 channels, which were colocalized (Fig. 1D-E). In contrast, GPR75-KO mice showed no signal for all channels, including GPR75 (Fig. 1F), indicating the specificity of GPR75 in situ probes. These results validate that GPR75-GFP-Ires-Cre mice express Cre and GFP in a faithful pattern to that of GPR75.

**Figure 1.**
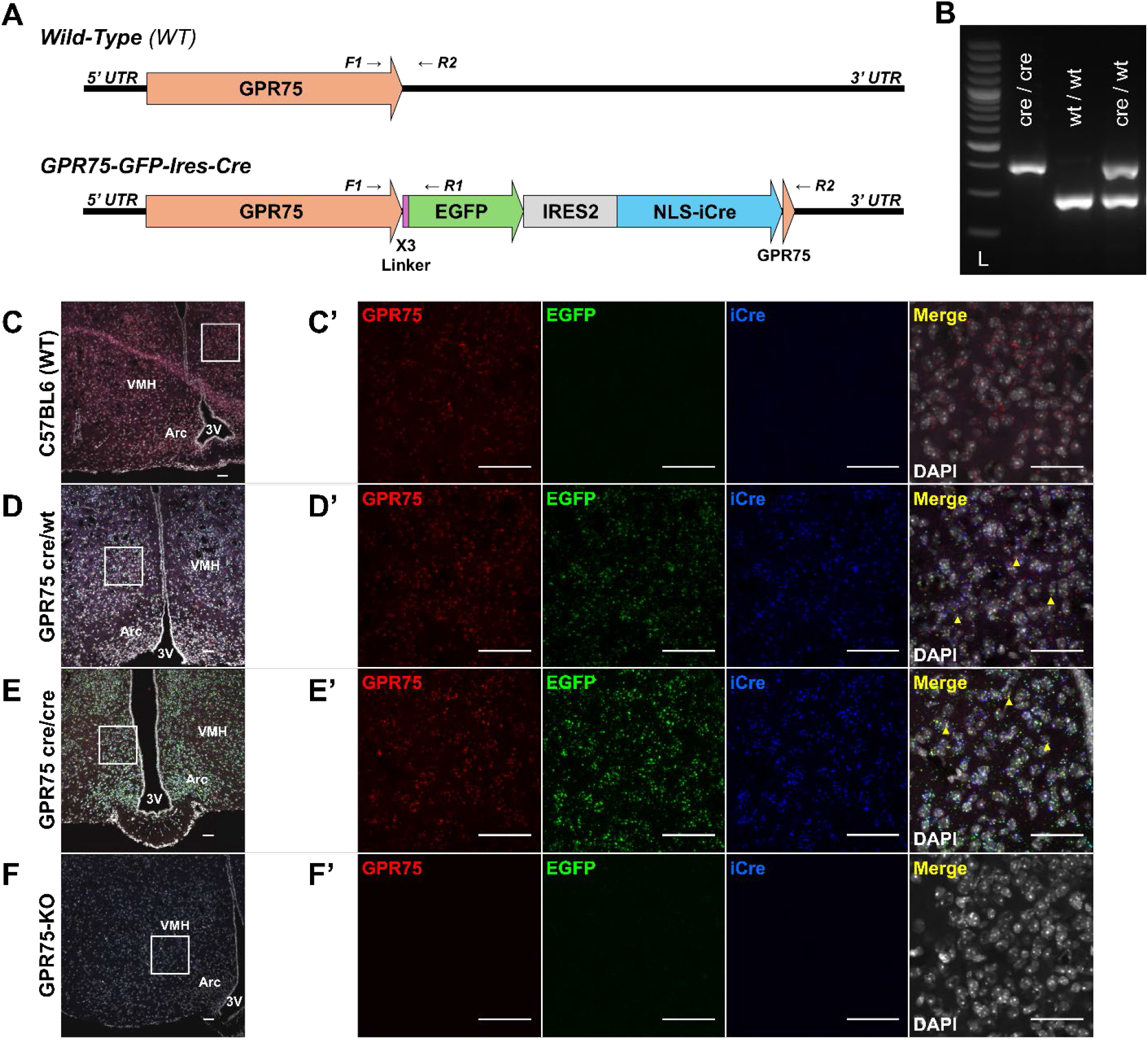
Development and Confirmation of Ires-Cre and EGFP Insertion into GPR75. **A**) Schematic of wild-type GPR75 gene construct and GPR75-GFP-Ires-Cre gene construct. **B**) Gel electrophoresis confirmation of Cre insertion, 100kB DNA ladder (lane 1; L), homozygous GPR75-GFP-Ires-Cre genotype (lane 2), Wild-type genotype (lane 3), heterozygous GPR75-GFP-Ires-Cre genotype (lane 4). C-F) In situ hybridization of GPR75 (red), EGFP (green), iCre (blue), and DAPI (white) for (**C**) wild-type, (**D**) heterozygous GPR75-GFP-Ires-Cre, (**E**) homozygous GPR75-GFP-Ires-Cre, and (F) total body deletion of GPR75 in the ventromedial hypothalamus. **D’-E’**). A yellow triangle points to colocalization of GPR75, EGFP, and iCre. **C-F**) Images taken at 20x magnification, (**C’-F’**) taken at 20x magnification with 5x zoom. Scale bar is 50µm in all images.

To examine the distribution of GPR75-expressing neurons in the brain, GPR75-GFP-Ires-Cre mice were bred with Cre reporter Rosa-Cas9-GFP mice to generate GPR75-GFP-Ires-Cre::Rosa-Cas9-GFP mice. Without immunostaining for GFP, GPR75-GFP-Ires-Cre mice do not show any GFP fluorescence (Fig. 2B-H). Thus, direct visualization of GFP fluorescence revealed the expression map of GPR75-expressing neurons across the mouse brain pseudo-colored red (Figure 2A-H). The GFP-expressing neurons were particularly abundant in the Nucleus Accumbens (Fig. 2B’), Suprachiasmatic Nucleus (Figure 2C’), Somatosensory Cortices (Figure 2D’), Medial Habenular Nucleus (Fig. 2D’’), Ventromedial Hypothalamus (Fig. 2D’’’), Arcuate Nucleus (Fig. 2E’), Ventral Tegmental Area (Fig. 2F’), Interpeduncular Nuclei (Fig. 2G’), and Cerebellum (Fig. 2H’). These results suggest that GPR75 is broadly expressed throughout the brain.

**Figure 2.**
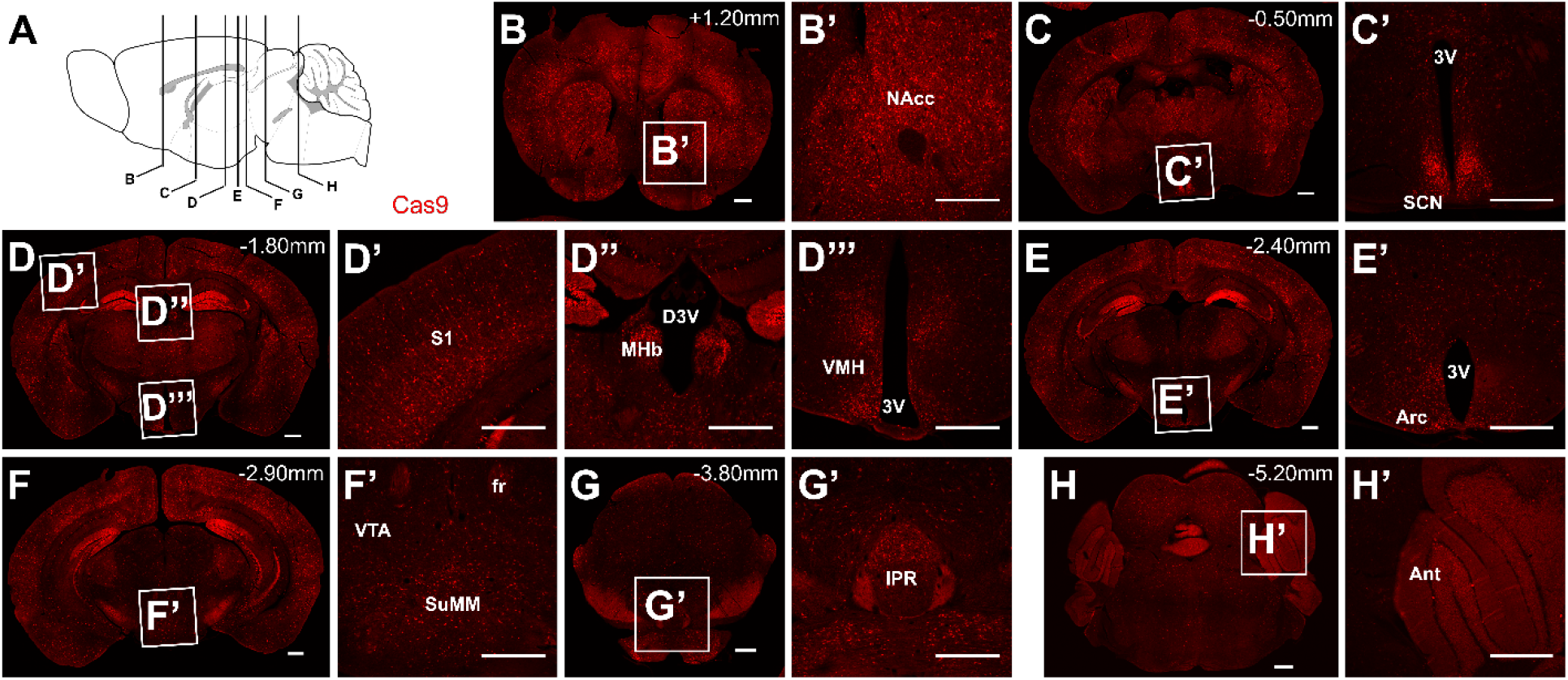
GPR75-GFP-Ires-Cre:Cas9 Whole Brain Expression. **A**) Schematic of slice locations on the sagittal mouse brain. Images are pseudo-colored red to distinguish between the endogenous GFP in figure 4. Stitched images taken at 10x of entire brain slices (**B, C, D, E, F, G, & H**). Individual images taken at 10x resolution (**B’, C’, D’, D’’, D’’’, E’, F’, G’, H’**). Bregma locations are at the bottom left of the whole slide images. Areas of interest include (B’) Nucleus Accumbens (NAcc), (C’) Third Ventricle (3V), Suprachiasmatic Nucleus (SCN), (D’) Primary Somatosensory Cortex (S1), (D’’) Medial Habenular Nucleus (MHb), Dorsal Third Ventricle (D3V), (D’’’) Ventromedial Hypothalamus (VMH), (E’) Arcuate Nucleus (Arc), (F’) Ventral Tegmental Area (VTA), Fasciculus Retroflexus (fr), Supramammillary Nucleus medial part (SuMM), (G’) Interpeduncular Nucleus rostral subnucleus (IPR), (H’) Anterior Lobe of the Cerebellum (Ant). Scale bar 500µm for all images.

Given the relatively large size of GFP, the potential fusion of GFP with GPR75 may disrupt its normal function. To this end, we examined body weight responses to HFD feeding in control wild-type mice and in heterozygous and homozygous GPR75-GFP-Ires-Cre mice. The mice were switched to HFD feeding after being reared on chow until 8 weeks of age. There was no difference in body weight among these mice during the period of 9 weeks when these mice were fed HFD in both males (Fig. 3A-C). In contrast, GPR75-KO mice exhibited a dramatic reduction compared to controls when fed HFD (Fig. 3D-F). These results collectively suggest that the normal function of GPR75 in the GFP fusion version remains intact.

**Figure 3.**
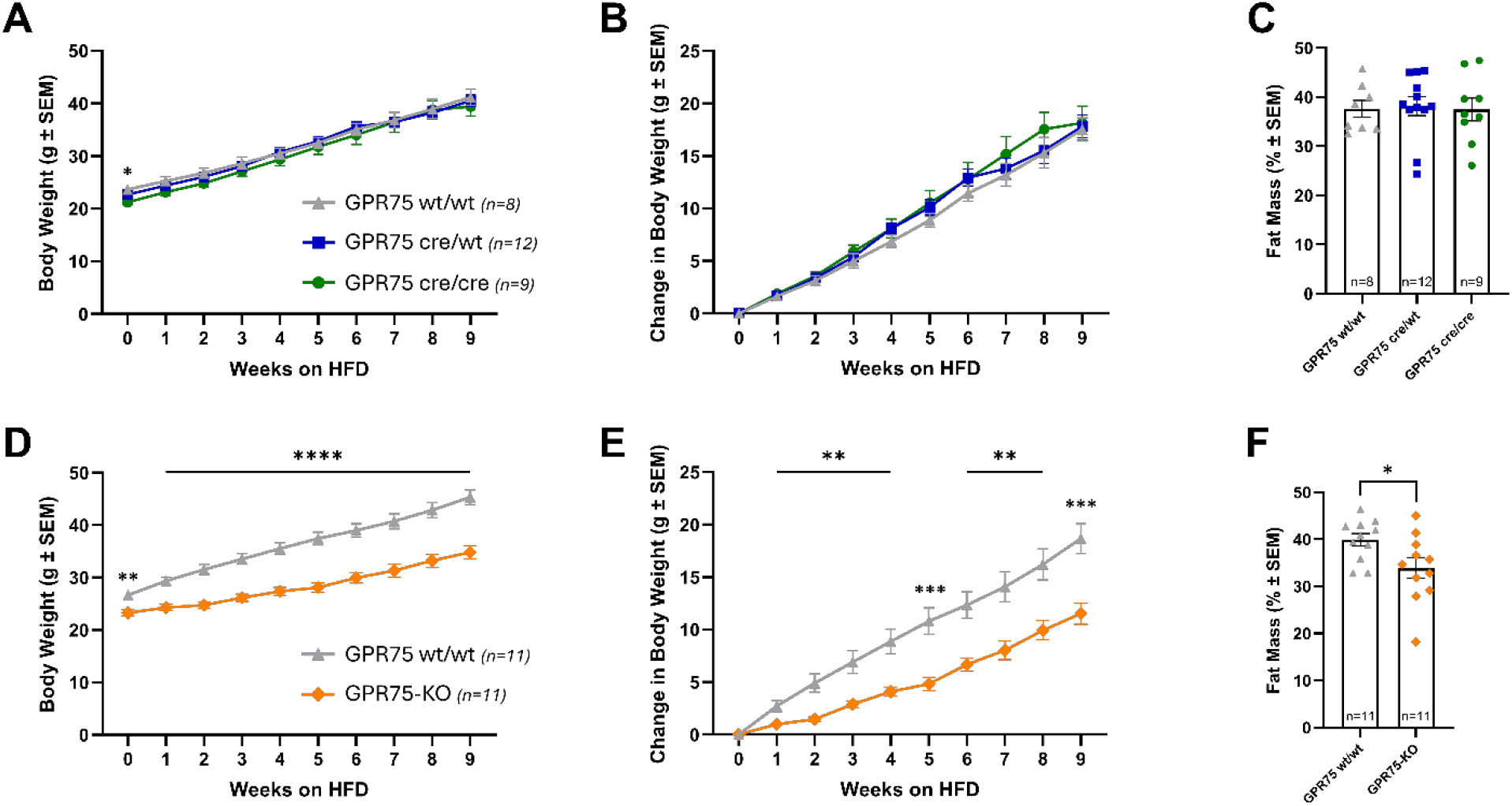
Insertion of EGFP and IRES-cre Does Not Alter Diet-Induced Obesity. **A-C**) Body weight and fat data for GPR75-GFP-Ires-Cre mice on HFD. **A**) Overall body weight gain on high fat diet in homozygous, heterozygous, and wild-type GPR75-GFP-Ires-Cre mice over 10 weeks (Two-Way Repeated Measures ANOVA, Week; F_(1.220, 31.72)_ = 356.9, p < 0.0001, Genotype; F_(2, 26)_ = 0.4474, p = 0.6441). **B**) Absolute body weight gain on HFD (Two-Way Repeated Measures ANOVA, Week; F_(1.220, 31.72)_ = 356.9, p < 0.0001, Genotype; F_(2, 23.96)_ = 0.4542, p = 0.6399). **C**) Percentage of fat mass to total mass after 10 weeks of HFD (One-Way ANOVA; F_(x,x)_ = x, P = x) . **D-F**) A separate cohort of mice, GPR75-KO and wild-type control were also fed HFD. **D**) Overall body weight gain on HFD in wild-type and GPR75-KO mice (Two-Way Repeated Measures ANOVA, Week; F_(1.272, 25.43)_ = 173.9, p < 0.0001, Genotype; F_(1, 20)_ = 36.55, p < 0.0001). **E**) Absolute body weight gain on HFD (Two-Way Repeated Measures ANOVA, Week; F_(1.231, 24.61)_ = 173.9, p < 0.0001, Genotype; F_(1, 20)_ = 17.27, p = 0.0005). **F**) Percentage of fat mass to total mass after 10 weeks of HFD (unpaired two-tailed t-test Welch Corrected; t_(16.58)_ = 2.356, P = 0.0311).

The *in vivo* intracellular localization of GPR75 remains unknown. Previous studies using a primary cell culture preparation suggest that the receptor is located in cilia ^10^. In GPR75-GFP-Ires-Cre mice, in which GFP was fused to GPR75, we failed to detect visible GFP fluorescence, likely due to very low *in vivo* expression of GPR75 protein. Detection with GFP antibodies revealed punctate GFP expression signal in the cytosol (Fig. 4B-C), which is specific, as no signaling was detected in wild-type control sections (Fig. 4A). Three-dimensional renderings of Z-stacks displayed the location of EGFP around the nucleus of neurons (Fig. 4D).

**Figure 4.**
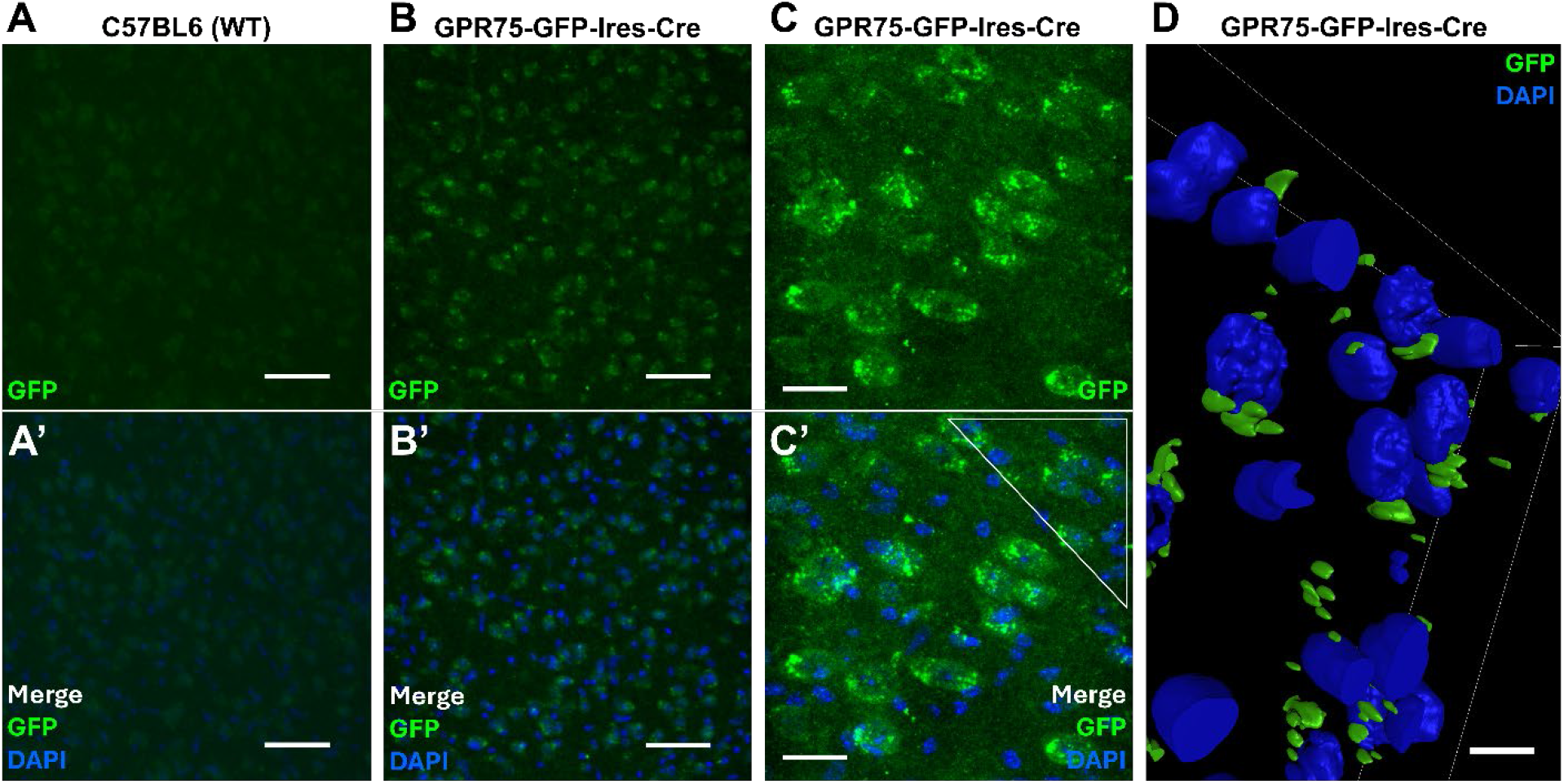
Cellular localization of GPR75 by GFP immunostaining in mouse brain tissue. **A-B)** Maximum intensity projection images of DAPI and GFP immunostaining acquired from an 11.02µm Z-stack (0.92µm Z-step) in C57BL6 (**A-A’**) and from a 12.04µm Z-stack (0.93µm Z-step) in GPR75-GFP-Ires-Cre (**B-B’**) mouse thalamic brain tissue taken at 20x magnification with a 3x zoom, scale bar is 50µm. **C-C’**) Maximum intensity projection image of GFP and DAPI immunostaining in GPR75-GFP-Ires-Cre mouse thalamic tissue acquired from a 16.44µm Z-stack (0.2µm Z-step) taken at 60x magnification with a 2x zoom, scale bar is 25µm. **D**) Representative 3D rendering of DAPI and GFP in GPR75-GFP-Ires-Cre mouse thalamic tissue. 3D rendering was compiled from a 16.44µm Z-stack (0.2µm Z-step), scale bar is 25µm.

## Discussion

Here, we generated and validated a new GPR75-GFP-Ires-Cre mouse line. Our data confirmed that the mRNA expression profiles of GPR75, GFP, and Cre are colocalized within the same neurons, as predicted from the knockin construct. We found that GPR75 is broadly expressed throughout all brain areas, consistent with other reports ^10,11^. Given this broad expression pattern, it is interesting that selective deletion of GPR75 in glutamatergic, but not GABAergic, neurons recaptures the resistance to DIO observed in GPR75-KO mice ^8^. Of note, previous studies suggest that glutamatergic, but not GABAergic, neurons mediate GLP-1R agonism in reducing obesity ^12^, demonstrating the specific function of glutamatergic neurons in mediating resistance to DIO.

Despite the known importance of brain GPR75 action in mediating DIO, its expression pattern in the brain remains obscure. We have tried several commercially available antibodies against GPR75 for immunostaining, but all have failed, despite the broad GPR75 mRNA expression pattern in the brain, suggesting very low GPR75 expression in the brain. Of note, previous studies using a strain of mouse with a GPR75-tag didn’t show any data on GPR75 expression in brain tissues, either, but instead showed data from a primary cell culture preparation^10^, likely due to difficulty in visualizing the endogenous GPR75 in brain tissues. For the current GPR75-GFP-Ires-Cre mice, we used a fresh 4% paraformaldehyde solution to fix the brain tissue, enabling the detection of GFP signal.

Intriguingly, our results with GFP immunostaining showed that GPR75 is mostly located in the cytosol, which is in contrast with the previous results in cell culture, suggesting that GPR75 is located in cilia ^10^. The reason for the observed difference in cellular localization is unknown, but it may be due to differences between *in vivo* brain tissues and *in vitro* cell culture. In summary, we have generated and validated a new GPR75-GFP-Ires-Cre mouse strain, which will allow for targeting GPR75 and GPR75-expressing neurons to study GPR75 function in mediating DIO.

## Materials & Methods

### Animals

All mice were group housed on a 12:12 light cycle and reared on standard rodent chow ad libitum until experimentation or tissue collection at approximately 8 weeks of age. For high-fat feeding studies, wild-type control and homozygous or heterozygous GPR75-GFP-Ires-Cre mice were switched to a high-fat diet (D12492, Research Diets Inc., New Brunswick, NJ) at 8 weeks of age for an additional 9 weeks. A second, separate cohort of GPR75 wild-type and GPR75 whole body KO (GPR75-KO) were fed HFD for 9 weeks. Rosa26-floxed STOP-Cas9-GFP mice (aka Rosa-Cas9-GFP) JAX stock #026175 were obtained from the Jackson Laboratory, and the GPR75^flox/flox^ strain ^13^ was generously provided by Dr. Darryl Zeldin at the National Institutes of Health. Weekly body weights were collected. The GPR75-KO allele was generated through germline deletion of the floxed GPR75 allele. Fat and lean mass data were collected using Echo MRI (Houston, TX). All experiments were approved by the IACUC of the University of Texas Health Science Center at Houston.

### Generation of GPR75-GFP-Ires-Cre Mice

The GPR75-GFP-Ires-Cre mouse strain was generated by Biocytogen (Waltham, MA, USA) using their Extreme Genome Editing System (EGE), a CRISPR/Cas9-based technology. Specifically, the 3XGGGGS-EGFP-Ires-Cre sequence was inserted after exon 2 of the GPR75 gene, replacing its stop codon. In this knockin strategy, GFP is fused to GPR75 via a 3XGGGGS linker, avoiding potential structural interference between GFP and GPR75, and the Ires sequence allows separate protein expression of GPR75-GFP and Cre from the same transcripts.

### Genotyping

Genotype confirmation of all mice was conducted using ear tissue. Tissue was placed in 50mM sodium hydroxide (SS276-1, Fisher Scientific) and heated at 98°C on a heat block for 45 minutes. An equal volume of 100mM Tris-hydrochloride was added to the mixture, which was then centrifuged for 10 minutes at 15,000 rpm. Primer solution was made using 5µL of 2x Taq Red Master Mix with 1.5mM MgCl (#5200300, Apex) 0.1µL of each primer, with ddH^2^O to a 10µL total volume per DNA sample. Samples were run on a 3% Agarose gel with a 100bp DNA ladder (#SM0241, GeneRuler, Thermo Scientific) for 45 minutes at 250V (400mA). Primers used for PCR genotyping are WT F: 5’ – GGCAAACTTGCCTCCAGTGAACTCTT; MUT R 5’ – TAGCGGCTGAAGCACTGCACG; and WT R 5’ – AATGTCAAGAAACCATGAAGCCAACC, which give rise to a band for the wild-type allele at 254 bps and the knockin allele at 352 bps.

### Immunostaining

Mice were transcardially perfused with 0.9% saline and 4% paraformaldehyde (PFA). Brains were collected and stored in 4% PFA overnight, followed by 30% sucrose solution until sunk. Brain tissue for immunofluorescence was sliced at 30µm, stored at 4°C in a 0.1% sodium azide (#19038, Acros) 1X PBS solution until stained and mounted for confocal microscopy.

GPR75-GFP-Ires-Cre and C57BL6 wild-type mouse brains were immunostained under identical conditions using a floating tissue protocol with 1X PBS and 0.1% Triton-X 100 (BP151, Fisher Scientific) as the stock solution and the aforementioned GFP antibody at a 1:500 dilution (GTX26673, GeneTex Inc.). Primary antibody host species was goat, normal donkey serum (#017-000-121, Jackson ImmunoResearch Laboratories) was used for non-specific blocking, and Alexa Fluor 633 Donkey anti-Goat secondary (A21082, Invitrogen) was used for fluorescent marking.

### In situ hybridization

Brain tissues used for *in situ* hybridization were sliced at 12µm, directly mounted to slides, and stored at -80°C until use. *In situ* hybridization of brain sections from wild-type C57BL6, GPR75-KO, and GPR75-GFP-Ires-Cre mice was performed using the RNAscope Multiplex Fluorescent V2 Assay kit (323100, ACDBio) according to the manufacturer’s guidelines. GPR75 (318281, ACDBio), EGFP-O4-C2 (538851-C2, ACDBio), and iCre-C3 (423321-C3, ACDBio) probes were utilized in this study. TSA vivid fluorophores 520 (323271, ACDBio), 570 (323272, ACDBio), and 650 (323273, ACDBio) were used at a dilution of 1:1000.

### Microscopy

Brain tissue slices were imaged on a Nikon AX-R Confocal using the NIS Elements software (Version 6.20.02, Nikon Instruments Inc.). Z-stack images for 3D renderings were acquired at 60x magnification with 2x zoom and processed in Amira 3D (Version 2025.1.1, Thermo Fisher Scientific).

### Statistical Analysis

Weekly body weight and change in body weight data were analyzed by two-way repeated measures ANOVA with Tukey’s multiple comparison test (Fig. 3A-B, 3D-E). Percentage fat mass data were analyzed by Welch’s ANOVA and unpaired t-test with Welch’s Correction for multiple comparisons (Fig. 3C) and Welch’s unpaired t-test (Fig. 3F). All statistical analyses and graphs were generated using GraphPad Prism 10 (Version 10.4.2, GraphPad Software, LLC). * p<0.05, ** p<0.01, *** p<0.001, **** p<0.0001.

## Acknowledgements

We acknowledge the Tong lab members for their helpful discussion. This work is partially supported by a training fellowship from UTHealth Houston Center for Clinical and Translational Sciences T32 Program (Grant No. T32 TR004904, EW), NIH R01 DK136284, R01 DK 135212, R01 DK 131466, R01DK109934, and DOD HT94252310156 (QT), and the Welch Foundation AU-0042-20030616 (ZA). QT is the holder of the Cullen Chair in Molecular Medicine and Hans J. Eberhard MD, PhD, and Irma Gigli, MD Distinguished Chair in Immunology at McGovern Medical School.

## Author contribution

The experiments were mainly conducted by E.W. with help from R.Z. and S.M.F.; S.Z. N.Z, Z.A provided essential reagents and guidance. E.W. wrote the manuscript with significant input from Q. T.

